# MCGA: a multi-strategy conditional gene-based association framework integrating with isoform-level expression profiles reveals new susceptible and druggable candidate genes of schizophrenia

**DOI:** 10.1101/2021.06.08.447608

**Authors:** Xiangyi Li, Lin Jiang, Chao Xue, Mulin Jun Li, Miaoxin Li

## Abstract

Linkage disequilibrium and disease-associated variants in non-coding regions make it difficult to distinguish truly associated genes from redundantly associated genes for complex diseases. In this study, we proposed a new conditional gene-based framework called MCGA that leveraged an improved effective chi-squared statistic to control the type I error rates and remove the redundant associations. MCGA initially integrated two conventional strategies to map genetic variants to genes, i.e., mapping a variant to its physically nearby gene and mapping a variant to a gene if the variant is a gene-level expression quantitative trait locus (eQTL) of the gene. We further performed a simulation study and demonstrated that the isoform-level eQTL was more powerful than the gene-level eQTL in the association analysis. Then the third strategy, i.e., mapping a variant to a gene if the variant is an isoform-level eQTL of the gene, was also integrated with MCGA. We applied MCGA to predict the potential susceptibility genes of schizophrenia and found that the potential susceptibility genes identified by MCGA were enriched with many neuronal or synaptic signaling-related terms in the Gene Ontology knowledgebase and antipsychotics-gene interaction terms in the drug-gene interaction database (DGIdb). More importantly, nine susceptibility genes were the target genes of multiple approved antipsychotics in DrugBank. Comparing the susceptibility genes identified by the above three strategies implied that strategy based on isoform-level eQTL could be an important supplement for the other two strategies and help predict more candidate susceptibility isoforms and genes for complex diseases in a multi-tissue context.

## Introduction

Genome-wide association studies (GWASs) have been used to identify novel genotype-phenotype associations for more than a decade, and thousands of single-nucleotide polymorphisms (SNPs) have been revealed for their associations with hundreds if not thousands of complex human diseases^1, 2^. Nevertheless, conventional GWAS analyses have limited power to produce a complete set of susceptibility variants of complex diseases^3^. Because most susceptibility SNPs only have small effects on a complex phenotype, conventional SNP-based association tests are generally underpowered to reveal susceptibility variants after multiple testing corrections. Moreover, the susceptibility variants scattering randomly throughout the genome are often in strong linkage disequilibrium (LD) with numerous neutral SNPs, and makes the discrimination of truly causal variants from GWAS hits quite difficult^3^. Finally, more than 90% of the disease-associated variants are in non-coding regions of the genome, and many of them are far from the nearest known gene, and it remains a challenge to link genes and a complex phenotype through the non-coding variants ^4, 5^. Accordingly, corresponding methodological strategies have been proposed to make up, at least in part, for the issues mentioned above.

First, gene-based approaches can reduce the multiple testing burden by considering the association between a phenotype and all variants within a gene^6^. Assigning a variant to a gene according to the physical position of the variant from gene boundary is one of the most popular strategies for gene-based approaches. For example, MAGMA (Multi-marker Analysis of GenoMic Annotation), one of the most popular gene-based approaches, uses a multiple regression approach to incorporate LD between markers and detect multi-markers effects to perform gene-based analysis^7^. VEGAS, a versatile gene-based test for GWAS, incorporates information from a full set of markers (or a defined subset) within a gene and accounts for LD between markers by simulations from the multivariate normal distribution^8^. GATES, a rapid gene-based association test that uses an extended Simes procedure to assess the statistical significance of gene-level associations ^9^.

Second, evaluating associations at one gene conditioning on other genes can isolate true susceptibility genes from non-susceptibility genes^10^. Yang et al. proposed an approximate conditional and joint association analysis based on linear regression analysis for estimating the individual causal variant with GWAS summary statistics^11^. Our previously proposed conditional gene-based association approach based on effective chi-squared statistics (ECS) could remove redundantly associated genes based on the GWAS p-values of variants. The comparison of ECS with MAGMA and VEGAS suggested that ECS might be more powerful to predict biologically sensible susceptibility genes ^10^.

Third, the observation that variants in the non-coding regions were enriched in the transcriptional regulatory regions implied that these variants might affect the disease risk by altering the genetic regulation of target genes^2^. Integration of expression quantitative trait loci (eQTL) studies and GWAS has been used to investigate the genetic regulatory effects on complex diseases. As many complex diseases manifested themselves in certain tissues, using the eQTLs of potentially phenotype-associated tissues might help identify the true susceptibility genes in tissue context^12^. Based on MAGMA, a method called eMAGMA, which integrated genetic and transcriptomic information (e.g., eQTLs) in a tissue-specific analysis to identify risk genes, was proposed to identify novel genes underlying major depression disorder^13^. S-PrediXcan was developed for imputing the genetically regulated gene expression component based on GWAS summary statistics and transcriptome prediction models built from the eQTL/sQTL dataset of the Genotype Tissue Expression (GTEx) project^14^. Researchers have applied S-PrediXcan to study genetic mechanisms of multiple complex traits ^15–17^. In contrast to the considerable research focusing on integrating gene-level eQTLs with GWAS summary statistics, little attention has been paid to integrating isoform-level expression QTLs (isoform-level eQTLs) with GWAS summary statistics. Michael J. Ganda et al. estimated the candidate risk genes of three psychiatric disorders based on GWAS summary statistics and isoform-level expression profiles. They emphasized the importance of isoform-level gene regulatory mechanisms in defining cell type and disease specificity^18^, and similar analyses and conclusions were generated for Alzheimer’s disease^19^.

Though much achievement has been attained, identifying independently phenotype-associated genes with high reliability remains challenging, especially for complex diseases. In the present study, we aimed to build a more powerful conditional gene-based framework based on a new ECS and investigate whether QTLs in phenotype-associated tissues, especially the isoform-level eQTLs, can predict more susceptibility genes or not. The main assumption of our study is that isoform-level eQTLs may reflect the more real variant-expression regulatory relationship than gene-level eQTLs, and using isoform-level eQTLs can help predict novel susceptibility genes and isoforms that cannot be found by the conventional gene-based approaches and gene-level eQTLs strategy. The formation procedure of the assumption is this: gene-level eQTLs are predicted based on gene-level expression profiles and corresponding genotype data. In contrast, isoform-level eQTLs are predicted based on isoform-level (or transcript-level) expression profiles and corresponding genotype data. The gene-level expression profiles are computed by averaging the expression of multiple isoforms belonging to the gene, which may omit the expression heterogeneity among these isoforms and neutralize the opposite effects and lower the power of gene-level eQTLs. Taken together, conventional gene-based approaches mainly focus on variants close to genes boundary (say +/-5kilo base pairs), thus omit remote but important association relationship between genes and variants. In the present study, we expanded the application scope of conventional gene-based approaches by using gene-level eQTLs and isoform-level eQTLs in the potentially phenotype-associated tissues to identify more candidate susceptibility genes and isoforms.

## Results

### Overview of the Multi-strategy Conditional Gene-based Association (MCGA) framework

Our previously powerful and unified framework, DESE, proposed to estimate the potentially phenotype-associated tissues of complex diseases, could iteratively run the conditional gene-based association analysis with the selective expression score of genes among multiple tissues^20^. We had demonstrated in DESE that the iterative operations with the selective expression analysis based on expression profiles could strengthen the power of conditional gene-based analysis. In this study, we proposed a new conditional gene-based framework, MCGA, which could also iteratively run conditional gene-based association analysis with selective expression analysis, to systematically explore the susceptibility genes associated with a complex phenotype using GWAS summary statistics and eQTL summary statistics of SNPs. MCGA has two main advantages over DESE. First, MCGA is based on a new effective chi-squared statistic (ECS), with which the type I error could be controlled within a proper level. Second, MCGA can perform conditional gene-based association analysis using different SNPs sets, i.e., physically nearby SNPs, gene-level eQTLs and isoform-level eQTLs.

To evaluate the performance of MCGA, we performed extensive simulations and a real data application to schizophrenia. Specifically, we organized the present study in four sequential parts that cover the optimizing the exponent of chi-squared statistics to control type I error rates, applying the improved chi-squared statistics to perform conditional gene-based association analysis in simulation data, applying the improved conditional gene-based association analysis to predict the potential susceptibility genes of schizophrenia, and finally extending the application by using gene-level eQTLs and isoform-level eQTLs. Three strategies were implemented in three conditional gene-based models, respectively. The model assigning a SNP to a gene according to the physical distance of the SNP from the gene boundary is named MCGA_Dist. The model using gene-level eQTLs is named MCGA_eQTL. The model using isoform-level eQTLs is named MCGA_isoQTL (**Figure 1**). All three models of MCGA have been implemented in our integrative platform KGGSEE.

**Figure 1:**
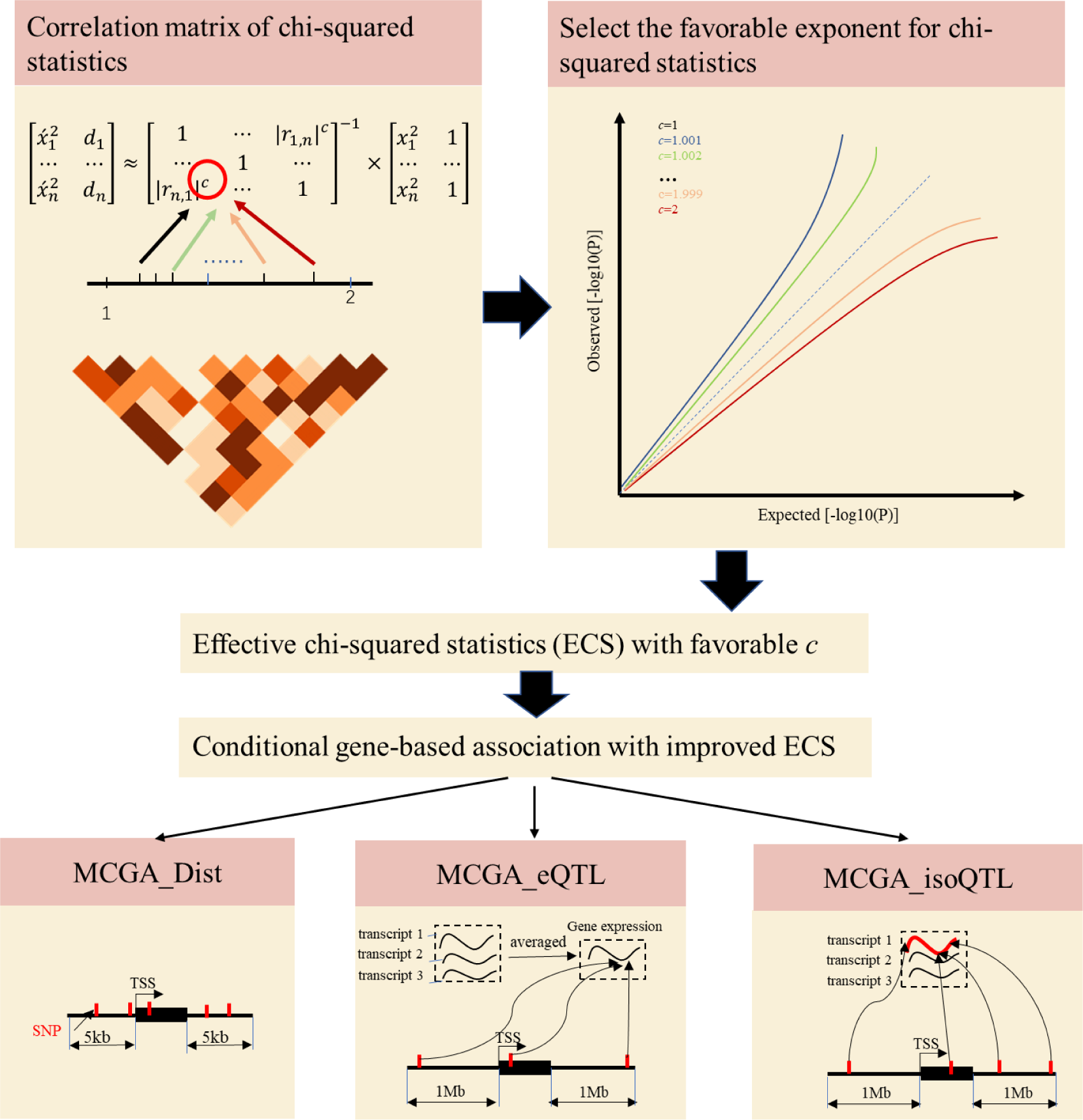
The simplified principle of the present study. First, choose the best exponent *c* between 1 and 2. Each time we increased *c* by an interval of 0.01. The best *c* can control the type I error within a proper level. Then we implemented the best *c* to improve the previous ECS approach and got the improved conditional gene-based approach. Third, the mapping strategies used by three conditional gene-based association models of MCGA. 5kb: 5000 base pairs. 1Mb: 10^6^ base pairs. TSS: Transcription Start Site.

Our simulation results showed that the type I error rate was controlled within a reasonable level by using the new effective chi-squared statistics (ECS) with the favorable exponent. Another simulation study pointed out that association analysis based on isoform-level eQTLs was more powerful than gene-level eQTLs. As for predicting the potential susceptibility genes of schizophrenia, MCGA_Dist, MCGA_eQTL and MCGA_isoQTL all produced a set of potentially susceptible and druggable genes. Moreover, with the help of MCGA_isoQTL, we also predicted the potential susceptibility isoforms (or transcripts) for schizophrenia. Our results also showed that the usage of isoform-level eQTLs could predict some important susceptible and druggable genes which cannot be found by MCGA_Dist and MCGA_eQTL.

### The favorable exponent *c* in the correlation matrix of chi-squared statistics to control the type I error rates

We found that the exponent *c* in the correlation matrix of chi-squared statistics could determine the divergence of the *p*-values of the ECS tests from the uniform distribution (see details in **Formula (1)** of Methods). Small *c* led to a p-value larger than expected, indicating the deflated type I error rate. Large *c* led to a p-value smaller than expected, indicating an inflated type I error rate. As shown in **Figure 2**, the *c*=1.0 led to deflated *p*-values while the *c*=2.0 led to inflated *p*-values in the upper tail of the Q-Q plot against the uniform distribution. This pattern was independent of sample sizes, variant sizes and phenotype distribution (binary or continuous) (**Figure 2**). The stable trend determined by the *c* value also implied that the favorable *c*, which could properly control the type I error rate, measured by the minimal mean log fold change (MLFC), must be within sections 1 and 2. Our theoretical derivation also demonstrated *c* value should be within section 1 and 2 (see details in the **Materials and Methods**). Moreover, it seemed given the *c* value, the distributions of *p*-values were similar at different sample sizes and phenotype distributions. As shown in the Q-Q plot (**Figure 3**), most majority *p*-values at the sample size 10,000 and 40,000 of binary or continuous phenotypes were overlapped. **Figure 3** showed the favorable c obtained by the grid search algorithm at 84 different scenarios. Again, the favorable *c* values were approximately independent of trait types, sample sizes and variant sizes. For the sake of simplicity, we proposed to use the averaged favorable *c* values, 1.432, for all the analyses in the present study.

**Figure 2:**
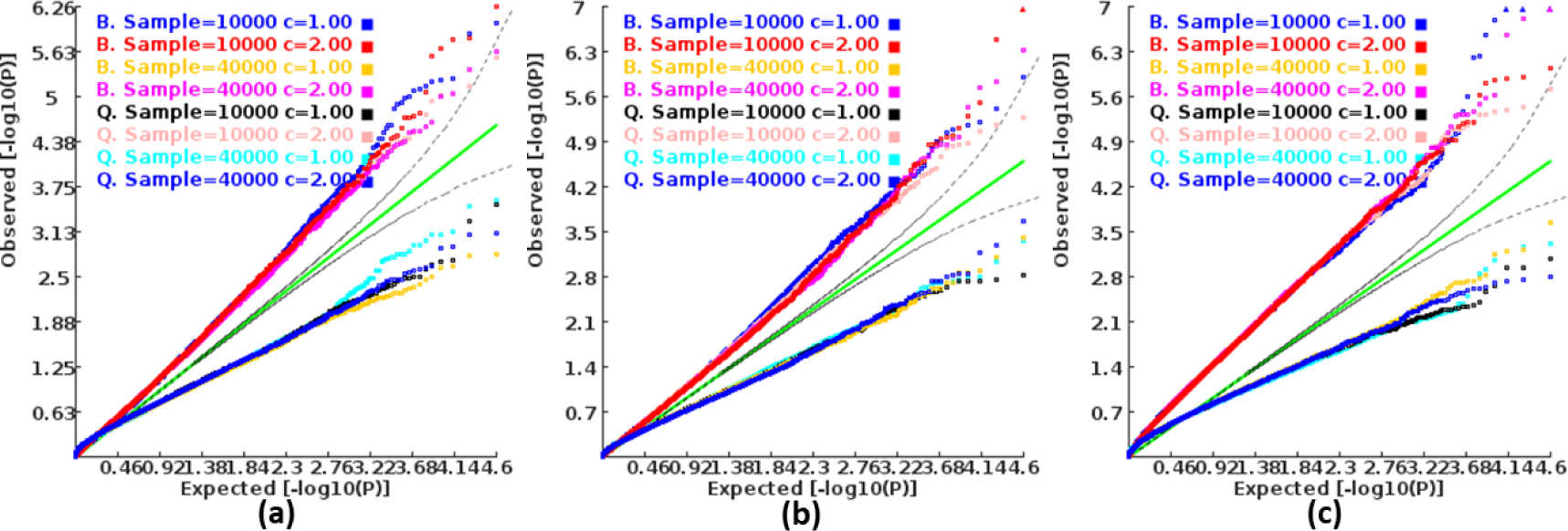
Q-Q plots of the *p*-value of the ECS test under null hypothesis based on two extreme exponents (i.e., 1 and 2). **a)**, **b)**, and **c)** represent the variant size of 50, 100 and 500, respectively.

**Figure 3:**
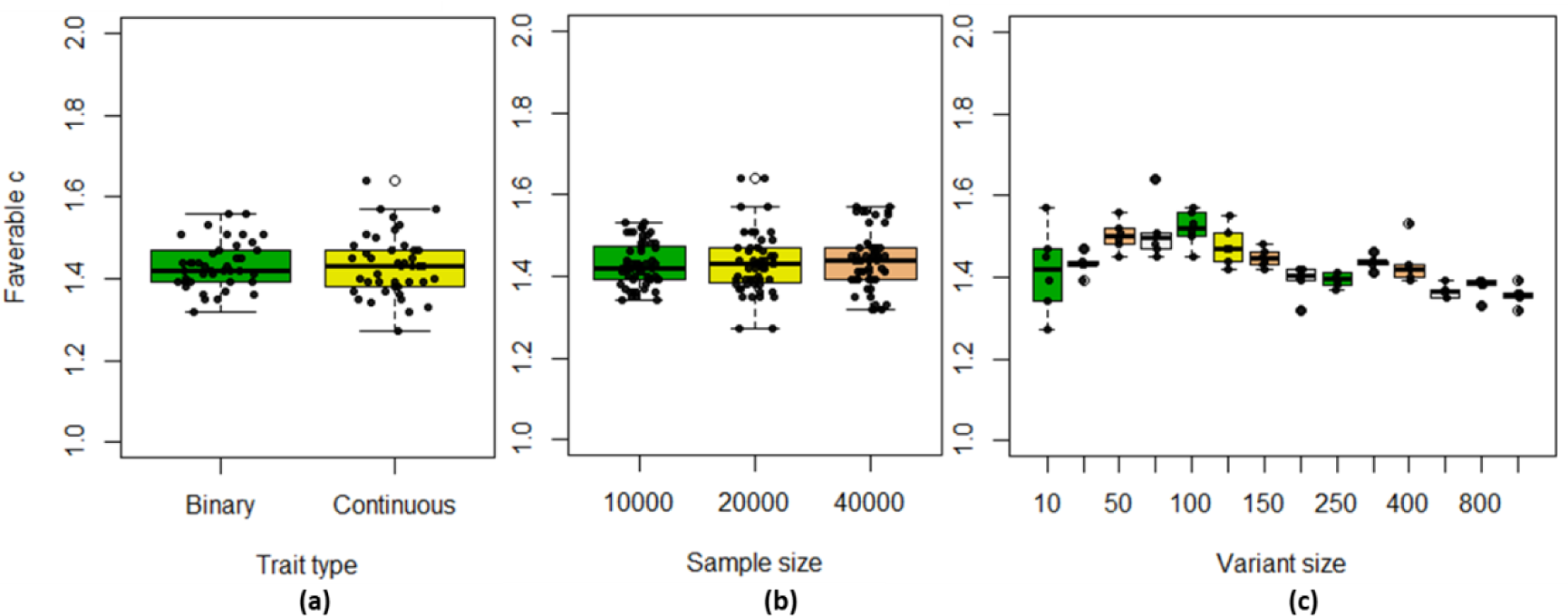
The boxplot of favorable *c* values at different simulation scenarios. **a)** at binary and continuous phenotypes; **b)** at different sample sizes; **c)** at different variant sizes.

### The type I error and power of 193 the conditional gene-based association analysis based on the new chi-squared statistics

Further, we investigated the type I error and power of the conditional gene-based association analysis based on the improved ECS with the above favorable exponent *c* value (i.e., 1.432). The basic function of the conditional gene-based test based on the improved ECS is to remove redundant associations among associated genes. As shown in **Figure 4**, in six different scenarios (see details in **Materials and Methods**), the conditional *p*-values of the genes without truly casual loci approximately followed the uniform distribution *U*[0,1] regardless of the variance explained by its nearby genes. The distribution of conditional *p*-value was similar to that produced by the conventional likelihood ratio test for nested linear regression models. These results suggested that the conditional gene-based association analysis based on the improved ECS could produce valid *p*-values for statistical inference. In contrast, the unconditional association test produced an inflated *p*-value due to the indirect associations produced by nearby causal genes in the LD block. Concerning the statistical power, we found conditional gene-based association analysis based on the improved ECS produced smaller *p*-values than the likelihood ratio test (**Figure 5**), suggesting a higher statistical power of the former. Another advantage of conditional gene-based association analysis based on the improved ECS over the likelihood ratio test was that the former did not require individual genotypes. The reason might be that the degree of freedom in the latter was inflated by the LD among variants. Hereinafter we named the conditional gene-based association analysis based on the improved ECS with favorable exponent *c* value as MCGA.

**Figure 4:**
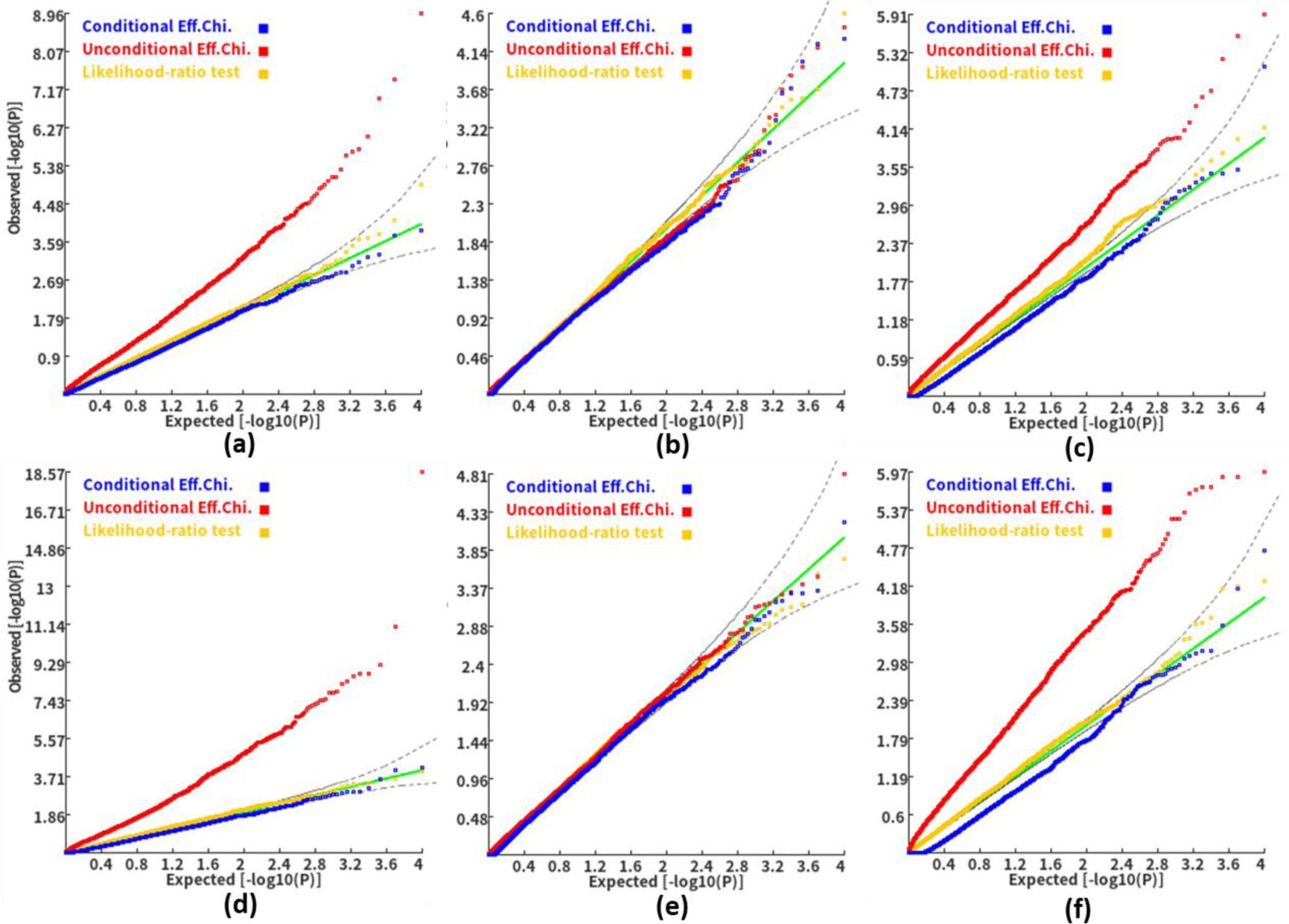
Q-Q plot of the conditional, unconditional gene-based association test and likelihood-ratio test under the null hypothesis. **a)** and **d)**: two gene-variant pairs with similar variant sizes (SIPA1L2 with 29 variants and LOC729336 with 30 variants). **b)** and **e)**: two gene-variant pairs with different variant sizes, and the first is larger than the second (CACHD1 with 41 variants and RAVER2 with 8 variants). **c)** and **f)**: two gene-variant pairs with different variant sizes, and the second is larger than the first (LOC647132 with 5 variants and FAM5C with 48 variants). **a)**, **b)** and **c)**: the former gene has no QTL, and QTL in the latter gene explained 0.5% of heritability. **d)**, **e)** and **f)**: the former gene has no QTL, and QTL in the latter gene explained 1% of heritability. Ten thousand phenotype datasets were simulated for each scenario. Unconditional Eff. Chi. (the red) represents unconditional association analysis at the former gene by the improved ECS. Conditional Eff. Chi (the blue) represents conditional association analysis at the former gene conditioning on the latter gene by the improved ECS. The likelihood ratio test (the yellow) was conducted based on nested linear regression models.

**Figure 5:**
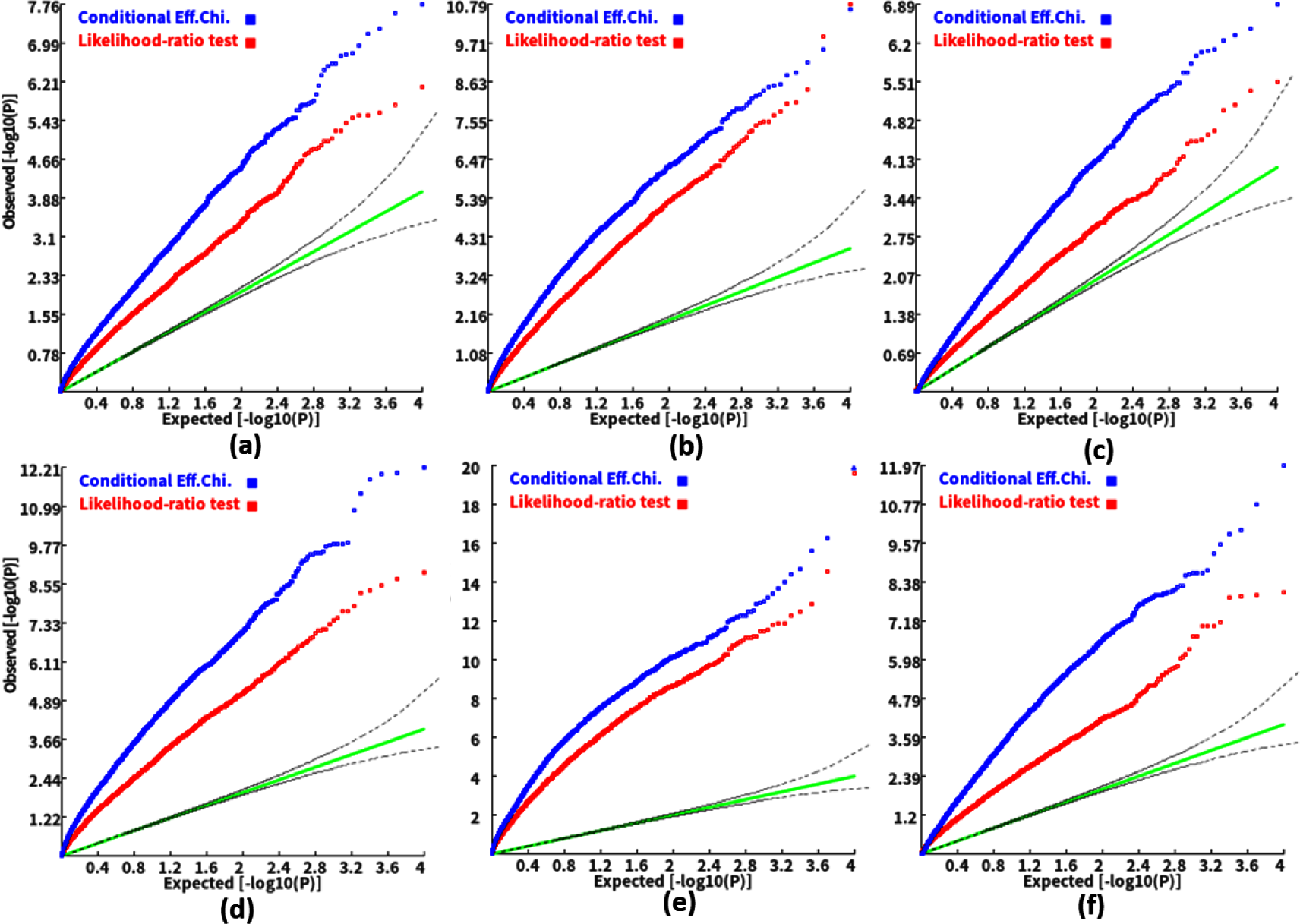
Q-Q plot of the conditional gene-based association test and likelihood-ratio test at different representative gene-variant pairs. **a)** and **d)**: two gene-variant pairs with similar variant sizes (SIPA1L2 with 29 variants and LOC729336 with 30 variants). **b)** and **e)**: two gene-variant pairs with different variant sizes, and the first is larger than the second (CACHD1 with 41 variants and RAVER2 with 8 variants). **c)** and **f)**: two gene-variant pairs with different variant sizes, and the second is larger than the first (LOC647132 with 5 variants and FAM5C with 48 variants). **a)-c)**: the QTL in either gene explained 0.25% of heritability. **d)-f)**: the QTL in either gene explained 0.5% of heritability. 1000 phenotype datasets were simulated for each scenario. Conditional Eff. Chi.: conditional association analysis at the former gene conditioning on the latter gene by the improved ECS. Likelihood Ratio Test: likelihood ratio test in which the full model included QTLs of both genes and the nested model included QTL of the latter gene.

### Application of MCGA to predict the potential susceptibility genes for schizophrenia (MCGA_Dist)

In the above simulation study, we demonstrated that the conditional gene-based analysis based on the improved ECS was more powerful than the likelihood ratio test in each simulation scenario. Here to further evaluate the performance of MCGA in the real-world data, we used a recent large-scale GWAS summary statistics dataset^20^ and gene expression profiles (GTEx v8) of ∼ 50 human tissues^21^ to identify the susceptibility genes of schizophrenia. Here, MCGA_Dist was first used, i.e., the variants were assigned to genes if the variants were in the window of +/-5kb around the gene boundary (see details in **Materials and Methods**). We found that 221 of 34,159 genes identified by MCGA_Dist had significantly statistical *p*-values smaller than 2.5E-6 (see details in **Table S1**).

**Table 1:** Different simulation scenarios and corresponding power in association analysis.

| Parameter combination |  | Power |  |  |  |
| --- | --- | --- | --- | --- | --- |
| Ev<sup>a</sup> | Eg | AllVar | isoform eQTL | eQTL_3isoform | eQTL_6isoform |
| 0 | 0.1 | 0 | **0.01** | 0 | 0.01 |
| 0.1 | 0.1 | 0.03 | **0.05** | 0.02 | 0.01 |
| 0.15 | 0.1 | 0.03 | **0.25** | 0.04 | 0.02 |
| 0.05 | 0.2 | 0.26 | **0.39** | 0.06 | 0.01 |
| 0.1 | 0.2 | 0.13 | **0.22** | 0.02 | 0.03 |
| 0.15 | 0.2 | 0.68 | **0.96** | 0.45 | 0.11 |

<sup>a</sup> **Ev** denotes the effect size of independent variants on gene expression. **Eg** denotes the effect size of gene expression on phenotype. More details can be seen in Methods.

To further study the functional annotations of the 221 potential susceptibility genes, we performed Gene Ontology (GO) enrichment analysis. Interestingly, we found that most GO:BP and GO:CC enrichment terms were neuronal-, dendrite- or synaptic signaling-related terms. Besides, the GO:MF enrichment terms were all about cellular signaling transduction (see examples in **Figure 6** and see details in **Table S2**). Systematic text-mining method was used to search the PubMed database to find papers that had reported the potential susceptibility genes of schizophrenia. The results showed that 87 of the 221 (∼ 39.4%) potential susceptibility genes had at least one search hit (see details in **Table S3**). The GO enrichment analysis and the text-mining results both implied the utility of MCGA_Dist.

**Figure 6:**
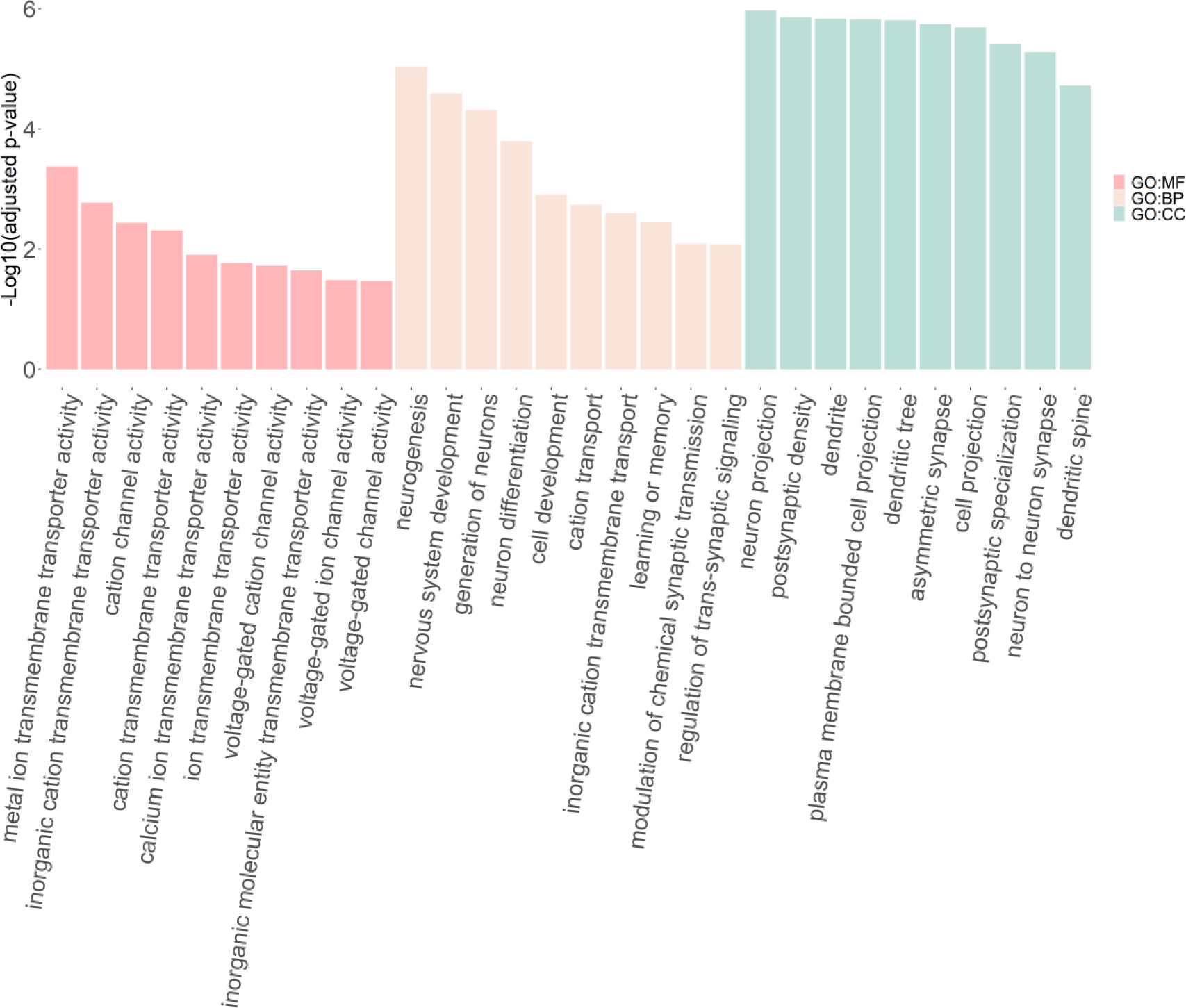
GO Functional annotations of the potential susceptibility genes of schizophrenia identified by MCGA_Dist. MF: Molecular Function of GO. BP: Biological Process terms of GO. CC: Cellular Component terms of GO. The X-axis represents the top 10 significant GO enrichment terms (in MF, BP and CC). The Y-axis represents the negative log10 of the adjusted *p*-value.

**Table 2:**
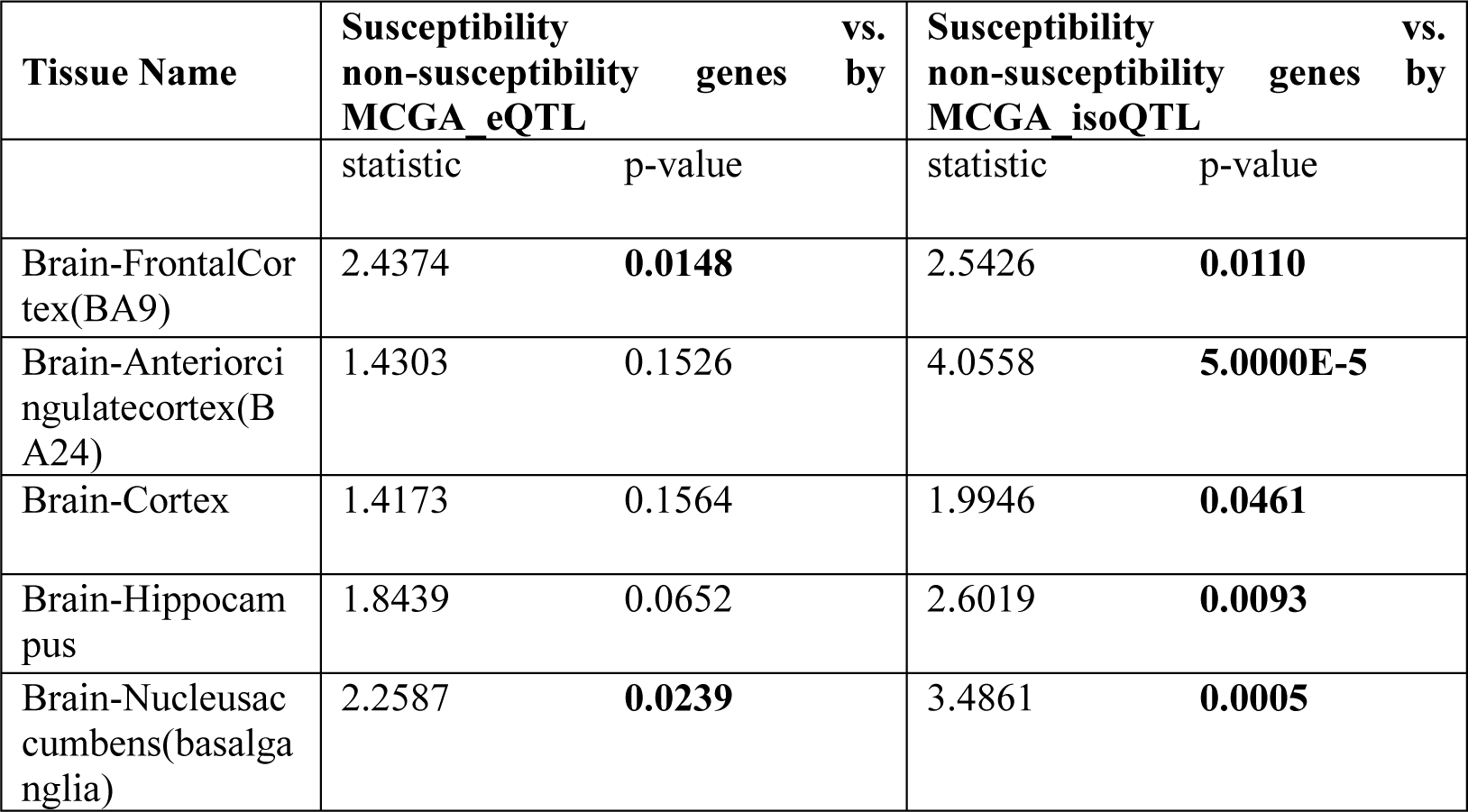
Comparison of the normalized intra-module connectivity of potential susceptibility genes identified by MCGA_eQTL and MCGA_isoQTL.

**Table 3:**
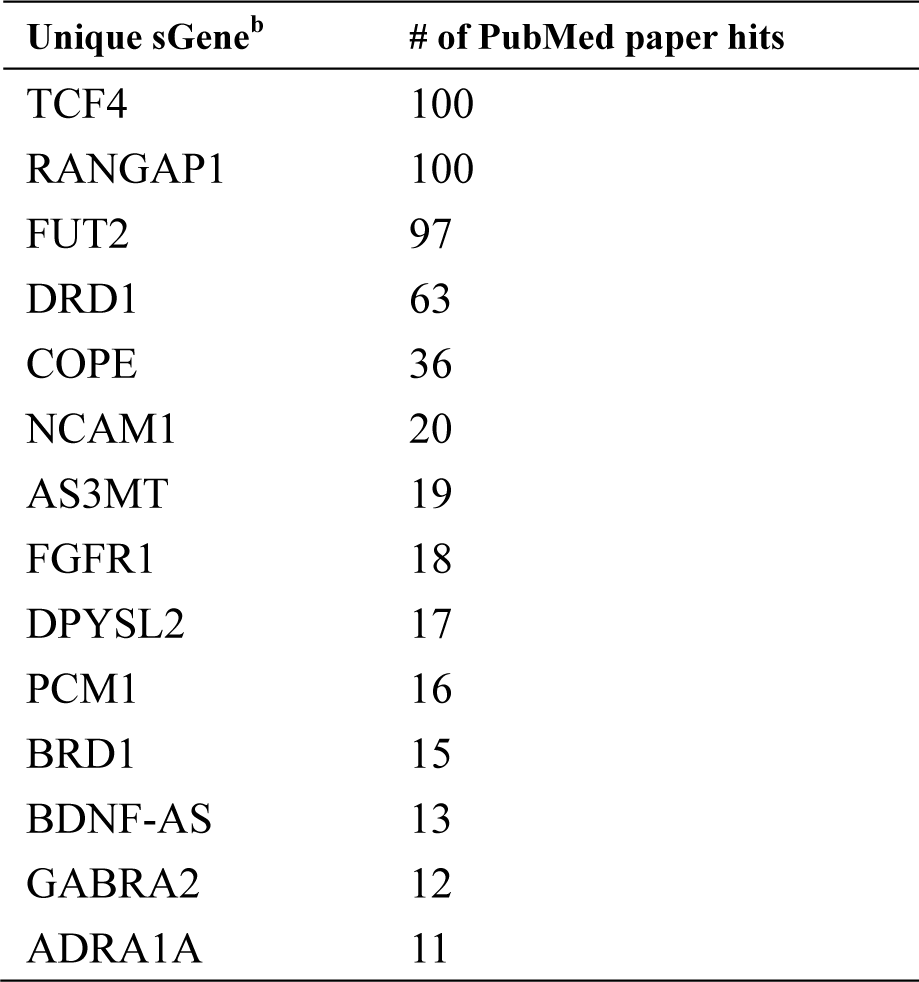
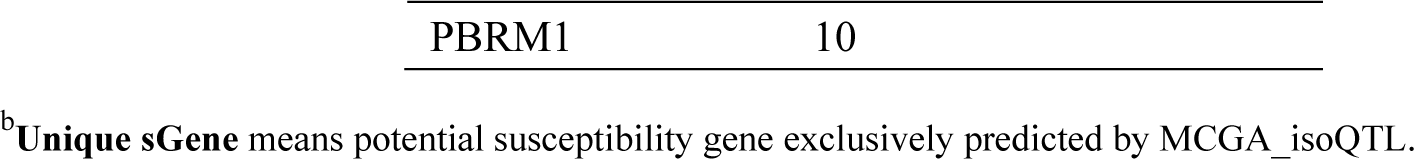
The important examples of potential susceptibility genes exclusively predicted by MCGA_isoQTL with at least ten search hits in PubMed.

### Evaluate the power of gene-level eQTLs and isoform-level eQTLs in association analysis by the simulation study

A common assumption is that genes close to significant variants are more likely to be the susceptibility genes, but the reality is that some potentially associated genes are not closest to the significant variants^16^. Molecular Quantitative Trait Loci (molQTL) is a genetic variant associated with a molecular trait, such as a gene-level eQTL and isoform-level eQTL, and can associate a variant with a gene or isoform. MCGA_Dist mapped a variant to a gene if the variant was in the +/-5kb window around the gene boundary. Next, we assigned a variant to a gene (or isoform) if the variant is a gene-level or isoform-level eQTL to broaden the application of MCGA. Since the isoform-level and corresponding gene-level expression profiles were quantified based on the same RNA-sequencing data, we wanted to test whether the power of association analysis based on the gene-level eQTLs was higher than that based on isoform-level eQTLs or not. We first performed a simulation study to evaluate the power of association analysis based on gene-level eQTLs and isoform-level eQTLs, respectively.

We considered the simplest case for simplicity, i.e., variants affected phenotype only through regulating the gene expression. We simulated genotype data, isoform-level expression profiles and corresponding phenotype data (see details in **Materials and Methods**). Specifically, we simulated four scenarios, i.e., association analysis using all variants (phenotype-associated isoform-level eQTLs and the other isoform-level eQTLs, denoted as Allvar in **Table 1**), association analysis only using phenotype-associated isoform-level eQTLs (denoted as isoform eQTL in **Table 1**), association analysis using gene-level eQTLs which were computed by the gene expression profiles derived by the average value of multiple isoforms belonging to the gene. As for scenarios of genes with multiple isoforms, we specifically simulated two new scenarios (denoted as eQTL_3isoform and eQTL_6isoform in **Table 1**), i.e., a gene with three (eQTL_3isoform) and six different isoforms (eQTL_6isoform). The expression value of the gene with three isoforms was averaged by the following three isoforms, i.e., one isoform associated with phenotype and the other two random isoforms simulated by the standard normal distribution *N*(0,1). The expression value of associated with phenotype and the other five random isoforms simulated by the standard normal distribution *N*(0,1) (see details in **Materials and Methods**). Based on the four scenarios mentioned above, we used six different parameter combinations to simulate six different cases, and each parameter combination was simulated 100 times to evaluate the statistical power. As shown in **Table 1**, the power of the association test based on phenotype-associated isoform-level eQTLs was the highest of all cases. The simulation results implied that isoform-level eQTLs were more powerful than gene-level eQTLs in association analysis.

### Broaden the application of MCGA by using gene-level eQTLs and isoform-level eQTLs (MCGA_eQTL and MCGA_isoQTL)

In the previous simulation study, we demonstrated that association analysis based on isoform-level eQTLs was more powerful than gene-level eQTLs in each simulation scenario. To further test this conclusion in real data and identify more potential susceptibility genes for schizophrenia, we first adopted the prioritized phenotype-associated tissues of schizophrenia by DESE^22^, a model to predict potentially phenotype-associated tissues based on gene selective expression analysis. The DESE results showed that all thirteen brain tissues were significantly associated with schizophrenia and ranked the top (**Figure 7a**). For the sake of simplicity, we then computed gene-level eQTLs of the top-five tissues based on gene-level expression profiles and isoform-level eQTLs of the top-five tissues based on transcript-level expression profiles, respectively. Hereinafter, the gene whose expression is associated with at least one SNP was denoted as eGene, and the gene with an isoform whose expression is associated with at least one SNP was denoted as sGene. Then we performed the improved conditional gene-based association analysis based on gene-level eQTLs and isoform-level eQTLs resulted from the corresponding tissues. In each of the top-five tissues, we found the number of potential susceptibility sGenes identified by MCGA_isoQTL was larger than that of potential susceptibility eGenes identified by MCGA_eQTL under the same filter cutoff 2.5E-6 (**Figure 7b**, see details in **Table S4** and **S5**). Besides, we found a considerable number of common genes between the estimated eGenes set and sGenes set in each of the top-five tissues (**Figure 7b**).

**Figure 7:**
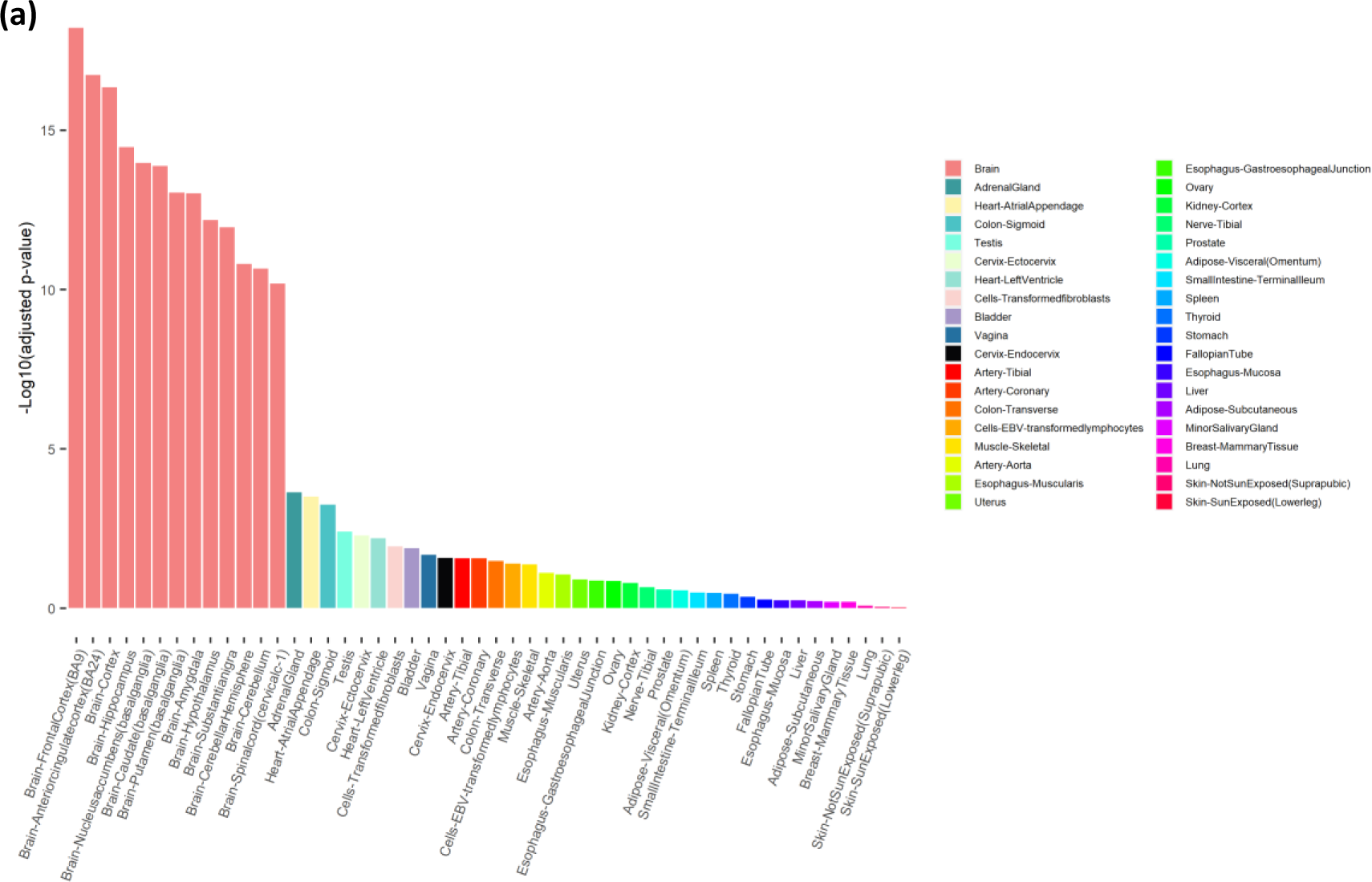
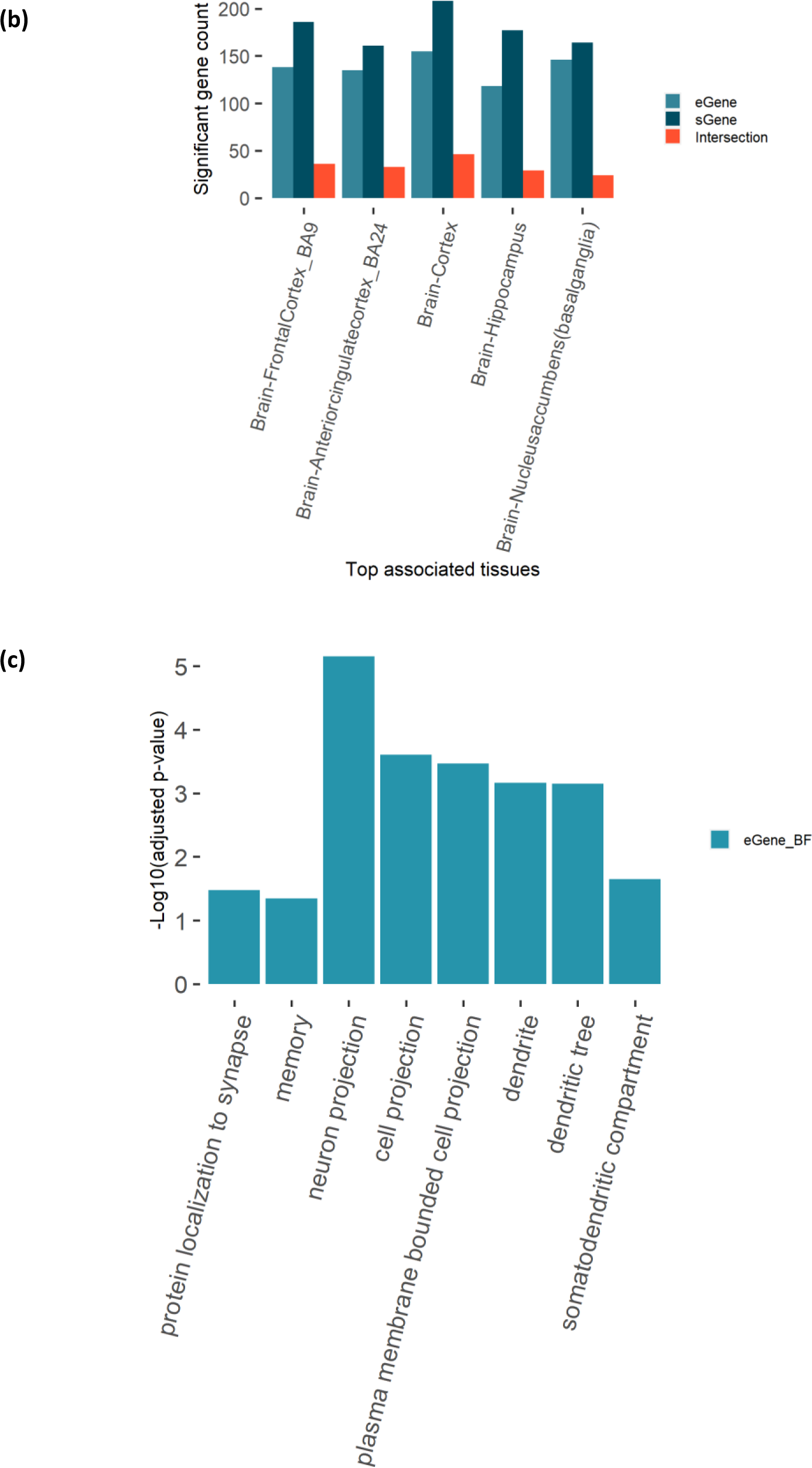
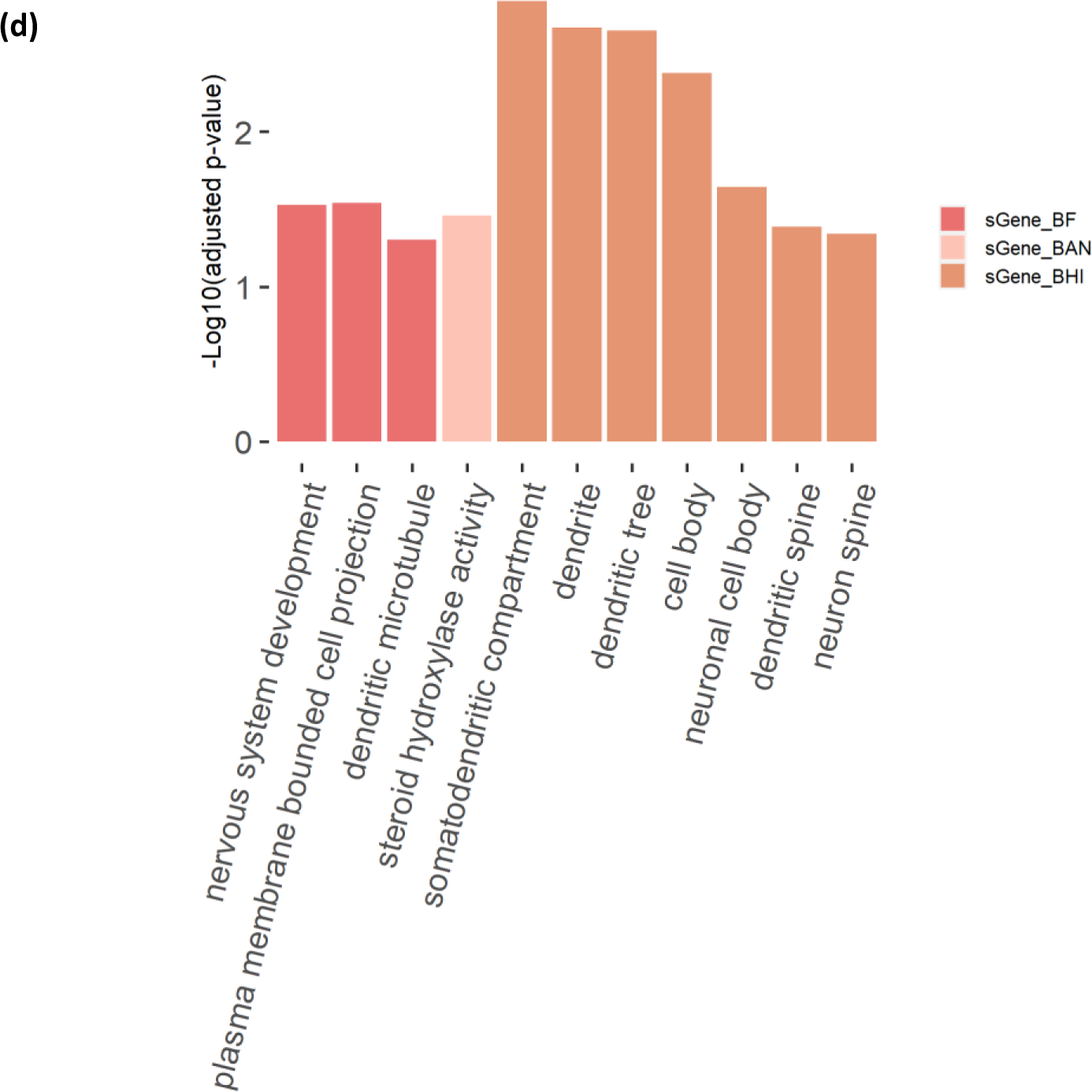
Potentially associated tissues of schizophrenia and the characterization of potential susceptibility genes in corresponding tissues. **a)** The statistical significance of tissues estimated to be associated with schizophrenia. The X-axis denotes the tissue names. The Y-axis means the negative log_10_(p-value) of the Wilcoxon rank-sum test. **b)** The comparison of eGenes and sGenes in each of the top-five associated tissues. **c**) The GO enrichment terms of eGenes in Brain-FrontalCortex(BA9). BF: Brain-FrontalCortex(BA9). **d**) The GO enrichment terms of sGenes in Brain-FrontalCortex(BA9), Brain-Anteriorcingulatecortex (BA24) and Brain-Hippocampus. BF: Brain-FrontalCortex(BA9). BAN: Brain-Anteriorcingulatecortex(BA24). BHI: Brain-Hippocampus.

**Table 4:** The potential susceptibility genes that are also the target genes of approved antipsychotics.

| Target gene | # of approved antipsychotics | Models |
| --- | --- | --- |
| DRD2 | 32 | MCGA_Dist & MCGA_eQTL |
| DRD1 | 22 | MCGA_isoQTL |
| ADRA1A | 21 | MCGA_isoQTL |
| CHRM3 | 10 | MCGA_eQTL |
| ADRA2B | 10 | MCGA_eQTL |
| HTR3A | 8 | MCGA_eQTL |
| OPRD1 | 1 | MCGA_Dist |
| GABRA2 | 1 | MCGA_isoQTL |
| CYP2D6 | 1 | MCGA_eQTL & MCGA_isoQTL |

We also performed the GO enrichment analysis to further investigate the functional annotations of these potential susceptibility eGenes and sGenes. For the eGene set in each of the top-five associated tissues, we found only the eGenes identified based on the gene-level eQTLs of Brain-FrontalCortex (BA9) had GO enrichment terms (**Figure 7c**). For the sGene set in each of the top-five associated tissues, we found the potential susceptibility sGenes in Brain-FrontalCortex(BA9), Brain-Anteriorcingulatecortex (BA24) and Brain-Hippocampus all had different GO enrichment terms (**Figure 7d**). We further combined the potential susceptibility eGenes and sGenes of all top-five tissues, respectively. Then we performed GO enrichment analysis based on the combined eGene set and sGene set. We found that both the eGene set and sGene set were enriched with many neuronal-, dendrite- or synaptic signaling-related GO terms (see details in **Table S6**). Then we searched the PubMed database with the combined eGene set (578 unique genes) and sGene set (696 unique genes), and found 133 of the 578 (∼ 23.0%) and 168 of the 696 (∼ 24.1%) potential susceptibility genes estimated by MCGA_eQTL and MCGA_isoQTL had at least one search hit, respectively (see details in **Table S7** and **S8**). The biologically sensible GO enrichment results and the PubMed search results both implied that the potential susceptibility sGenes and eGenes might have strong associations with schizophrenia.

### The advantages of MCGA_isoQTL versus MCGA_Dist and MCGA_eQTL

The connectivity score of a gene in the weighted co-expression network might imply its real association with other genes, and highly connected genes are often defined as hub genes. These hub genes are located in or near the center of corresponding co-expression modules and might play important roles in trait development^23^. We built an unsigned weighted co-expression network for each top-five tissue and investigated the normalized intra-module connectivity of potential susceptibility genes in each co-expression module. We found that the normalized intra-module connectivity scores of potential susceptibility genes in Brain-FrontalCortex_BA9 and Brain-Nucleusaccumbens(basal ganglia) identified by MCGA_eQTL were significantly larger than that of non-susceptibility genes. Interestingly, the normalized intra-module connectivity scores of potential susceptibility genes identified by MCGA_isoQTL were significantly larger than that of non-susceptibility genes in all the top-five schizophrenia-associated tissues (Wilcoxon rank-sum test p<0.05) (**Table 2**).

We next compared the potential susceptibility genes predicted by MCGA_Dist, MCGA_eQTL and MCGA_isoQTL. As shown in **Figure 8**, twenty-three genes were collectively predicted to be susceptible to schizophrenia by the three models of MCGA. As MCGA_isoQTL could output susceptibility gene-isoform pairs of corresponding tissues, we further got the corresponding susceptibility isoforms of twenty-three genes in corresponding tissues (see details in **Table S9**). Interestingly, we found that susceptibility isoforms for a gene varied greatly in different tissues. For example, *ABCC8* and *LINC01415* both had one susceptibility isoform in the top-five tissues. *ENST00000529967* of *ABCC8* was significantly associated with schizophrenia only in Brain-Hippocampus, while *ENST00000587320* of *LINC01415* was significantly associated with schizophrenia in Brain-Cortex, Brain-FrontalCortex(BA9) and Brain-Nucleusaccumbens(basal ganglia). We also found that different isoforms of the same gene were predicted to be significantly associated with schizophrenia in different tissues, such as *ENST00000377600* of *BTN2A1* significantly associated with schizophrenia in Brain-Cortex and *ENST00000312541* of *BTN2A1* significantly associated with schizophrenia in Brain-FrontalCortex(BA9). MCGA_isoQTL can help predict potential susceptibility genes and isoforms of corresponding phenotype-associated tissues at a more precise level.

**Figure 8:**
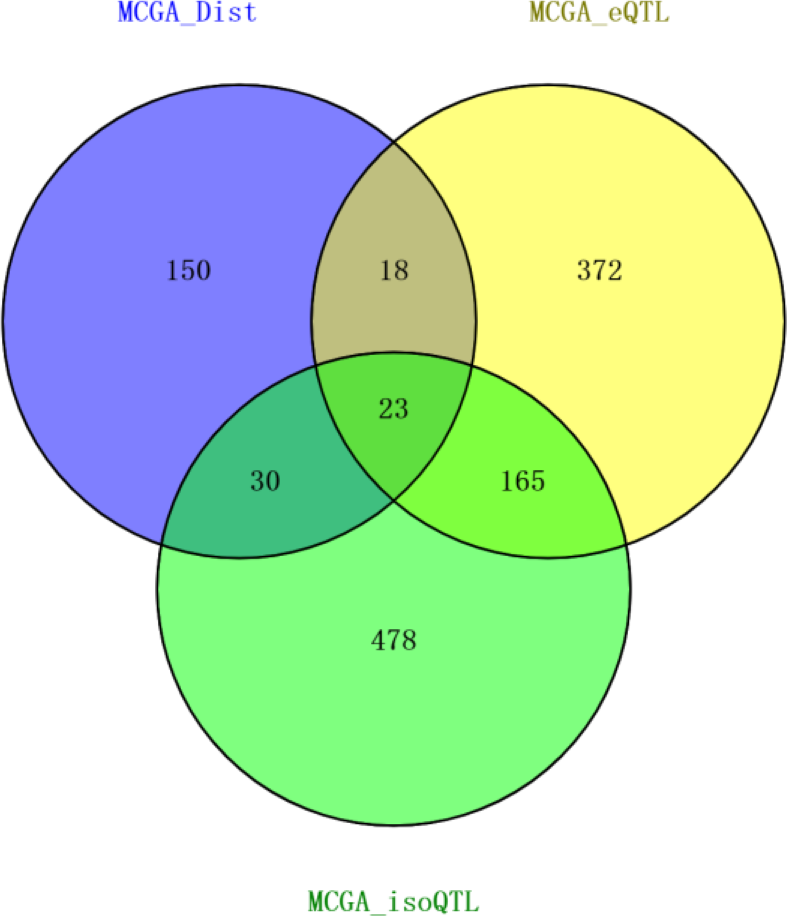
Comparison of the potential susceptibility genes predicted by MCGA_Dist, MCGA_eQTL and MCGA_isoQT. The venn plot was used to show the intersection genes and unique genes among MCGA_Dist, MCGA_eQTL and MCGA_isoQTL.

Except for the advantage of identifying the potential susceptibility isoforms, we found that some of the susceptibility genes exclusively predicted by MCGA_isoQTL were also biologically sensible. We found 478 susceptibility genes exclusively predicted by MCGA_isoQTL (**Figure 8**). We searched the PubMed database with these exclusive genes. The search results showed that 101 of the 478 (21.1%) genes each had a least one search hit which reported the associations of these exclusive genes with schizophrenia. Moreover, 15 of the 478 (3.1%) exclusive genes each had at least ten different supported papers in PubMed. Interestingly, transcription factor 4, i.e., *TCF4*, was reported by 100 papers in the PubMed database. *TCF4* is broadly expressed and may play an important role in nervous system development [provided by RefSeq, Jul 2016]. The important examples of biologically sensible genes exclusively identified by MCGA_isoQTL were listed in **Table 3**.

Taken together, from the perspective of WGCNA, the statistical significance of the comparison between potential susceptibility genes and non-susceptibility genes implied that these susceptibility genes identified based on isoform-level eQTLs might play more important roles in the weighted gene co-expression network of corresponding tissues. Our results also suggested that incorporating with isoform-level eQTLs can help predict more potential susceptibility genes than gene-level eQTLs in each potentially phenotype-associated tissue. Our results pointed that MCGA_isoQTL could help find some novel and important susceptibility genes which cannot be found by MCGA_Dist and MCGA_eQTL. Moreover, based on the isoform-level eQTLs of each phenotype-associated tissue, the MCGA_isoQTL strategy can also predict the potential susceptibility isoforms in the corresponding tissues.

### The druggability of the potential susceptibility genes identified by MCGA

Since drug target genes with genetic support are twice or as likely to be approved than target genes with no known genetic associations^24, 25^, we searched the DrugBank 5.0 database^26^ and found that nine potential susceptibility genes identified by MCGA were the target genes of multiple FDA-approved antipsychotics (**Table 4**). Several most popular target genes of approved antipsychotics, i.e., *DRD2*, *DRD1* and *ADRA1A*, were identified by different MCGA models and the results suggested that the three models could complement each other to identify more potential target genes.

To further investigate the druggability of the potential susceptibility genes, we searched the Drug Gene Interaction database (DGIdb v4.2.0) ^27^ and filtered the drug-gene interaction terms with at least one supported PubMed paper. After the filtration, we kept 30,072 unique drug-gene interaction terms and found 679 unique drug-gene interaction terms for 34 FDA-approved antipsychotics (see details in **Table S10**). Then we put the full list of potential susceptibility genes (by MCGA_Dist, MCGA_eQTL and MCGA_isoQTL, respectively) into DGIdb to investigate if the “antipsychotic”-“susceptibility gene” interactions were enriched in DGIdb. As shown in **Table 5**, we found that “antipsychotic” - “potential susceptibility genes” identified by the three models of MCGA were all significantly enriched in DGIdb. Moreover, as shown in **Figure 8**, 372 out of 578 and 478 out of 696 potential susceptibility genes were exclusively identified by MCGA_eQTL and MCGA_isoQTL, respectively. We found 253 unique drug-gene interaction terms for susceptibility genes exclusively predicted MCGA_eQTL (see details in **Table S11**), and 17 of 253 interaction terms were antipsychotics-gene interactions (hypergeometric distribution test *p*-value= 7.05E-5). We also found 291 unique drug-gene interaction terms for susceptibility genes exclusively predicted MCGA_isoQTL (see details in **Table S12**), and 28 of 291 interaction terms were antipsychotics-gene interactions (hypergeometric distribution test *p*-value = 1.48E-10). We also investigated the potential druggability of the susceptibility genes identified by MCGA. Among the 42 potentially druggable gene categories in DGIdb, we found the top three potentially druggable categories for the susceptibility genes identified by MCGA_Dist, MCGA_eQTL and MCGA_isoQTL were all “DRUGGABLE GENOME” (63 vs. 129 vs. 137. The number denotes the gene set size belonging to this category, same as below), “ENZYME” (33 vs. 68 vs. 90) and “KINASE” (22 vs. 40 vs. 60) (see details in **Table S13**). Taken together, our results showed that some of the potential susceptibility genes identified by MCGA had the potential to be druggable, and the application of eQTLs (especially the isoform-level eQTLs) could aid MCGA to identify more potentially druggable genes.

**Table 5:** The enrichment of drug-gene interaction terms in DGIdb for susceptibility genes identified by MCGA.

| Models | # of total drug-gene interaction terms | # of antipsychotics-gene interaction terms | Enrichment $p^c$ |
| --- | --- | --- | --- |
| MCGA_Dist | 332 | 33 | $1.37E-12$ |
| MCGA_eQTL | 831 | 64 | $1.39E-17$ |
| MCGA_isoQTL | 792 | 46 | $6.66E-9$ |

<sup>c</sup>Enrichment $p$ denotes the $p$ -value of hypergeometric distribution test.

## Discussion

In this study, we proposed a multi-strategy conditional gene-based association framework, MCGA, based on a new correlation matrix of chi-squared statistics to identify the potential susceptibility genes and isoforms for complex phenotypes. Comparing with the unconditional association test and likelihood ratio test, MCGA showed a lower type I error rate and higher statistical power. Since MCGA is a gene-based method, in this study, we adopted three strategies to map a variant to a gene, i.e., mapping based on physical position, gene-level eQTLs and isoform-level eQTLs. We implemented these three mapping strategies in corresponding three conditional gene-based association models, i.e., MCGA_Dist, MCGA_eQTL and MCGA_isoQTL, to predict the potential susceptibility genes for schizophrenia.

MCGA_Dist could output a list of genes, while MCGA_isoQTL and MCGA_eQTL could produce a list of genes for each potential phenotype-associated tissue because of the usage of gene-level eQTLs and isoform-level eQTLs of each tissue. Though MCGA_Dist predicted a relatively small size of susceptibility genes, these genes were enriched with plenty of neuronal- or synaptic signaling-related GO terms. Similar results of other research were obtained by gene-set analyses, which demonstrated that genetic variants associated with schizophrenia were enriched with synaptic pathways^28^. Besides, considerable amounts of these genes had been reported by many research papers in the PubMed database to support their associations with schizophrenia.

Since MCGA_Dist might omit some remote but important gene-variant associations, we improved MCGA_Dist with MCGA_eQTL and MCGA_isoQTL. We performed a simulation study and demonstrated that isoform-level eQTLs were more powerful than gene-level eQTLs in association analysis. Moreover, we found in real data that the size of the susceptibility gene set for schizophrenia predicted by MCGA_isoQTL was larger than MCGA_eQTL in each phenotype-associated tissue under the same threshold. Further, we found MCGA_isoQTL had two advantages over MCGA_eQTL and MCGA_Dist. First, several important potential susceptibility genes were exclusively predicted by MCGA_isoQTL. For example, fifteen potential susceptibility genes exclusively predicted by MCGA_isoQTL each had at least ten search hits in PubMed, which implied these genes were popular in schizophrenia studies. Second, to our best knowledge, MCGA_isoQTL was the first conditional gene-based association approach to produce a list of phenotype-associated isoforms (or transcripts).

In addition, we investigated the druggability of the susceptibility genes for schizophrenia identified by MCGA. Several susceptibility genes identified by MCGA were also the popular target genes of multiple FDA-approved antipsychotics. Besides, the “susceptibility gene”- “antipsychotics” interactions were enriched in DGIdb. The druggablilty of the important susceptibility genes, especially the sGenes identified based on isoform-level eQTLs, provided more credible supports for the utility of MCGA.

Our framework might have three potential applications. First, MCGA_Dist can be used to predict potential susceptibility genes and isoforms for other complex phenotypes. Second, based on the assumption that the distribution of expression profiles of true susceptibility genes might change before and after therapeutic drug treatment, MCGA_Dist can be used to perform drug repositioning analysis based on the drug perturbed expression profile. Third, since MCGA_eQTL and MCGA_isoQTL can help predict potential susceptibility genes in each potential phenotype-associated tissue, our framework can help perform synergistic drug combination prediction to screen drugs that can simultaneously perturb the expression of potential susceptibility genes in each potential phenotype-associated tissue.

The present study was limited by several factors. First, the moderate sample size (ranging from 129 ∼ 205) and mixed populations in GTEx v8 might both reduce the accuracy of gene/isoform-level eQTLs. Future genetic studies based on increased sample sizes might alleviate this problem. Second, the size of the susceptibility genes identified by MCGA_eQTL (578) and MCGA_isoQTL (696) was a little larger than conventional studies. One of the reasons might be that five brain regions were involved in the present study, and each brain region might have very different dysfunctional genes associated with schizophrenia. We also used MAGMA to identify the susceptibility genes of schizophrenia with the same GWAS summary statistics and found that MAGMA also identified ∼ 600 potential susceptibility genes with the basic parameter setup (see details in **Table S14**). Susceptibility genes identified by MCGA_eQTL and MCGA_isoQTL had many biologically meaningful annotations (such as neuronal- or synaptic signaling-related terms) in the GO databases, and some susceptibility genes were the target genes of multiple antipsychotics, and more than 20% of the susceptibility genes had been previously reported by other schizophrenia research in the PubMed database. Though these potential susceptibility genes were lack of systematically experimental validation, we shared the potential susceptibility genes in Table **S1**, **S4** and **S5** and encouraged follow-up studies to evaluate the function and roles of these susceptibility genes in the development of schizophrenia.

In conclusion, in this study, we proposed a new statistical framework to predict potential susceptibility genes for complex phenotypes based on GWAS summary statistics and gene/isoform-level eQTLs in a multi-tissue context. The application of our framework to schizophrenia revealed many novel susceptible and druggable genes. Besides, the usage of isoform-level eQTLs can be an important supplement for the conventional gene-based approach. The framework was packaged and implemented in our integrative platform KGGSEE (http://pmglab.top/kggsee/#/). We hope our framework can facilitate researchers to gain more insights into the phenotype-associated genes and isoforms of complex phenotypes.

## Materials and Methods

### The new effective chi-squared statistics (ECS) for conditional gene-based association analysis

We improved our previously proposed effective chi-squared test^10^ for a more efficient conditional gene-based association analysis based on a new correlation matrix of chi-squared statistics. The improved effective chi-squared statistics had two methodological advances to address the potential inflation issue, i.e., a type I error-controlled correlation matrix of the observed chi-squared statistics and a non-negative least square solution for the independent chi-squared statistics. The reasoning process was as follows. Suppose there were *n* loci in a set of genes. One wanted to calculate the association *p*-value of another physically nearby gene (containing *m* loci) conditioning on the set of genes (*n* loci). The first step of the conditional analysis was to produce effective chi-squared statistics for the set of genes (*n* loci) and all the genes (*n*+*m* loci in total).

Each locus had a *p*-value for phenotype association in the GWAS. The *p*-values were converted to corresponding chi-squared statistics with the degree of freedom 1. According to Li et al.^10^, each locus could be assumed to have a virtually independent chi-squared statistic. An observed marginal chi-squared statistic of a locus was equal to the summation of its virtually independent chi-squared statistic and the weighted virtually independent chi-squared statistic of nearby loci. The weight was related to the chi-squared statistics correlation, which was a key parameter of the analysis. The correlation of chi-squared statistics between two loci was approximated by the absolute value of genotypic correlation to the power of *c*, i.e., |*r*|^*c*^. Here, we derived that the key parameter, i.e., exponent *c*, ranged from 1 to 2, corresponding to different non-centrality parameters of a non-central chi-squared distribution (See the derivation in the next section). According to Li et al.^10^, the *n* virtually independent chi-squared statistics of the gene set could be approximated by a linear transformation of the *n* observed chi-squared statistics (**Formula (1)**),

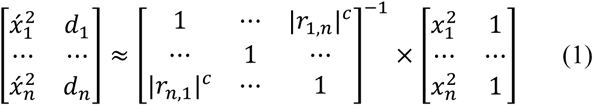

where 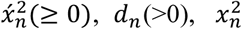 and |*r_i,j_*| denoted a virtually independent chi-squared statistic, degree of freedom of the virtually independent chi-squared statistic, an observed chi-squared statistic and the absolute value of the LD correlation coefficient (approximated by genotypic correlation), respectively. The effective chi-squared statistic 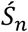 with the degree of freedom 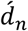 of the *n* loci was then obtained by **Formula (2)**:

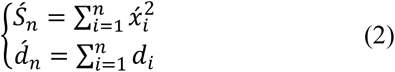

The effective chi-squared statistics 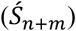 and degree of freedom 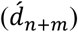 of the *n*+*m* loci could be calculated in the same way.

The effective chi-squared statistics of the *m* loci conditioning on the *n* loci was then approximated by **Formula (3)**,

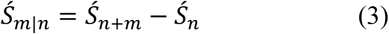

with the degree of freedom 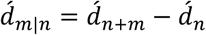.

Because 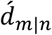 was no longer an integer, we used the Gamma distribution to calculate the *p*-values. Given the above statistics and degree of freedom, the *p*-value was equal to 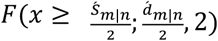, where the *F*(*x*) function was the cumulative distribution function of a Gamma distribution.

Because the virtually independent chi-squared statistics and degrees of freedom were expected to be larger than 0, we adopted a sequential coordinate-wise algorithm to approximate them^29^. This algorithm avoided unstable solutions in the above linear **Formula (1)** due to stochastic errors in the correlation matrix and observed chi-squared statistics.

After the above multiple approximations, it was still difficult to obtain the analytic solution for the exponent *c* in **Formula (1)**. We proposed a grid search algorithm to find a favorable value of exponent *c* to control type I error rates of the effective chi-squared tests. The error rate was examined by divergence from a uniform distribution between an obtained and theoretical top 1% *p*-values given a *c* value, measured as mean log fold change (MLFC) ^30^. In the grid search process, we increased *c* from 1.00 to 2.00 by an interval of 0.01 because it ranged from 1 to 2 (see the derivation in the **Materials and Methods**). The *c* value leading to the minimal MLFC was defined as the favorable *c* value. We considered in total 84 parameter settings, i.e. a combination of three different sample sizes (10,000, 20,000 and 40,000) and 14 different variant sizes (10, 30, 50, 80, 100, 125, 150, 200, 250, 300, 400, 500, 800, and 1000) for both binary and continuous traits, respectively. For a parameter setting, 40,000 datasets were simulated and used to produce *p*-values to determine the favorable *c* value for the setting. A region on chromosome 2 [chr2: 169428016-189671923] was randomly drawn for the simulation. The allele frequencies and LD structure of variants in the European panel of the 1000 Genomes Project were used as a reference to simulate genotype data by the HapSim algorithm ^31^. According to either the Bernoulli distribution or Gaussian distribution, each subject was randomly assigned a phenotype value under the null hypothesis. The Wald test under either logistic regression or linear regression in which the major and minor allele was encoded as 0 and 1 was used to produce the association *p*-value at each variant. The *p*-values of the variants were then analyzed by the effective chi-squared test for the gene-based association analysis.

### Approximate the correlation of chi-square statistics under the alternative hypothesis

Let two normal random variables 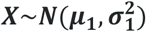 and 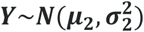 have covariance ***c***. Note that a squared normal random variable has non-central chi-square distribution and the squared mean of the former is called noncentrality parameter. The two variables can also be factorized as 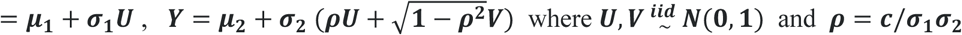.

Then we can calculate the co-variances of the two non-central chi-square variables ***X***^2^ and ***Y***^2^ by the factorized variables, 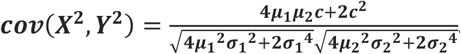. Suppose ***X*** is the Z score of the true casual variant and ***Y*** is the Z score of a non-functional variant in LD (coefficient ***r***) with the causal variant. One can assume 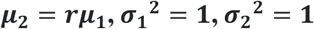 and ***c* = *r*.** Therefore, the correlation of ***X***^2^ and ***Y***^2^ can be simplified as, 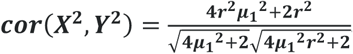.

Under the null hypothesis, ***μ***_1_ = 0 then ***cor***(***X*^2^, *Y*^2^**) = ***r***^2^. Under the alternative hypothesis of large scaled sample, the ***μ***_1_ or the noncentrality parameter becomes very large, correlation of ***X***^2^ and ***Y***^2^ become close to ***r***. ***μ*** → ∞, 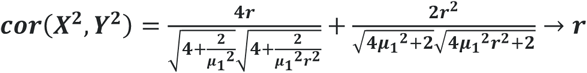.

Overall, the correlation between the two (non-central) chi-square ranges from *r*^2^ to *r*.

### The conditional gene-based association analysis for genome-wide association study

In a GWAS, all genes were firstly calculated with the *p*-values of unconditional gene-based association test using the above effective chi-squared statistics. For a given *p*-value cutoff, the significant genes were extracted and subjected to the conditional gene-based association analysis. When there were multiple significant genes in an LD block, the genes were conditioned one by one in a pre-defined order. In the present conditional analysis, the order of the gene was defined according to the unconditional *p*-value of the gene. Here we assigned the genes within 5 Mb into the same LD block. The conditional *p*-value of the first gene was defined as its unconditional *p*-value. The conditional *p*-value of the second gene was obtained by conditioning on the first gene, and that of the third gene was obtained by conditioning on the top two genes. The conditional *p*-values of subsequent genes were calculated according to the same procedure.

### Simulations for investigating type I error and power of the conditional gene-based association analysis

Extensively independent computer simulations based on a different reference population (i.e., EAS) in different genomic regions were performed to investigate type I error and the power of the conditional gene-based association test. To approach the association redundancy pattern in realistic scenarios, we used real genotypes and simulated phenotypes. The high-quality genotypes of 2,507 Chinese subjects from a GWAS were used ^32^, and phenotypes of subjects were simulated according to the genotypes under an additive model. Given total variance explained by *n* independent variants, V_g_, the effect of an allele at a bi-allelic variant was calculated by 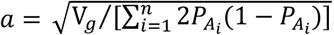, where *P_A__i_* was the frequency of alternative alleles. The total expected effect *A* of a subject was equal to a*[the number of alternative alleles of all the n variants]. Each subject’s phenotype was simulated by *P*=A+e, where *e* was sampled from a normal distribution *N*(*0*, 1-V_g_). We randomly sampled three pairs of genes, i.e., *SIPA1L2* vs. *LOC729336*, *CACHD1* vs. *RAVER2*, and *LOC647132* vs. *FAM5C*. The three pairs of genes represented three scenarios where the nearby gene (i.e., the first gene) had similar (*SIPA1L2* vs. *LOC729336*), larger (*CACHD1* vs. *RAVER2*) and smaller (*LOC647132* vs. *FAM5C*) variant size than the target gene (i.e., the second gene) in terms of SNP number, respectively. In the type I error investigation, the target gene had no QTLs, while the nearby gene had one or two QTLs. In the investigation of the statistical power, both genes had QTLs.

For power comparison, the likelihood ratio test based on linear regression was adopted to perform the conditional gene-based association analysis with raw genotypes. In the full model, genotypes of all SNPs encoded as 0, 1, or 2 according to the number of alternative variants entered the regression model as explanatory variables. In the subset model, the SNPs of the nearby genes entered the regression model. The calculation of the likelihood ratio test was performed according to the conventional procedure. The R packaged “lmtest” (version 0.9.37) was adopted to perform the likelihood ratio test.

### Simulations for comparing the power of gene-level eQTLs and isoform-level eQTLs in gene-based association tests

We compared the power of conventional gene-based association tests, gene-level eQTLs guided gene-based association tests and isoform-level eQTLs guided gene-based association tests by simulation studies. Assume some variants regulate gene expression, and the gene expression subsequently influences the phenotype. The same region on chromosome 2 [chr2: 169428016-189671923] was considered for the simulation. In the EUR panel of 1000 Genomes Project^33^, this region contains 1600 common variants (MAF>0.05). Genotypes of the variants were simulated given allelic frequencies and LD correlation matrix according to the HapSim algorithm^31^. Phenotypes were simulated under a polygenic model of random effect^34^. According to severe LD pruning (*r*^2^<0.01), eighty-two independent variants were extracted from the 1600 variants. The SNPs’ genotypes (*s*) contributing to the phenotypes were then standardized as, 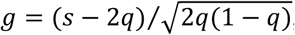, where *q* was the allele frequency of alterative allele. Phenotypes were simulated under a polygenic model of random effect^34^. We assumed 40% of the independent causal variants (*m*_X_) regulated gene expression (total heritability 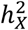). The expression of a gene (X) was simulated according to **Formula (4)**:

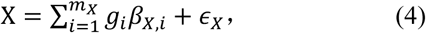

where 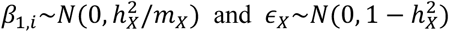.

The gene expression then contributed δ to a phenotype (Y). The phenotype value was simulated according to the **Formula (5)**:

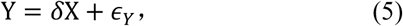

where *ϵ_Y_*∼*N*(0, 1 − δ^2^). Here Y was a continuous phenotype. For a binary phenotype, a cutoff *t* was set according to a given disease prevalence *K* under a standard normal distribution and the liability threshold model ^35^. Subjects with simulated Y values ≥*t* were set as patients, and others were set as normal controls.

When a gene had multiple isoforms, we assumed one of the isoforms was associated with phenotype and simulated expression values of the isoform according to the above regulation model (**Formula (5)**). The expression values of the remaining isoforms were simulated by the standard normal distribution *N*(0,1). The expression profile of a gene with multiple isoforms was averaged by the expression profiles of all the isoforms belonging to the gene. The gene-level eQTLs and isoform-level eQTLs were examined by the Wald test under the linear regression model. The variant-phenotype association analysis was performed based on the conventional association analysis procedure, and the statistical significance cutoff was set at *p*-value<0.001.

### Genome-wide association study of schizophrenia

The GWAS summary statistics of schizophrenia included 53,386 cases and 77,258 controls of European ancestry (hg19 assembly). Genotypes in the CEU panel from the 1000 Genomes Project were used to correct for the relatedness of the summary statistics. To predict the potential susceptibility genes of schizophrenia, the variants in the major histocompatibility complex (MHC) region, i.e., chr6:27,477,797-34,448,354, were excluded because of high polymorphism in the present study. Detailed descriptions of population cohorts, quality control methods and association analysis methods can be found in reference^20^. The summary statistics can be accessed at the Psychiatric Genomics Consortium.

### The Genotype-Tissue Expression (GTEx) project

The GTEx project (release v8) created a resource including whole-genome sequence data and RNA sequencing data from ∼ 900 deceased adult donors^21^. Four tissues or cell types (i.e., whole blood, cells-Leukemiacellline_CML, pancreas and pituitary) were filtered out and not included in the following analyses because of the small sample size or weak correlation of gene expression profiles with most of the other tissues.

### GO annotation of the potential susceptibility genes

Functional enrichment analyses were performed by g:Profiler^36^. GO terms, i.e., biological processes (BP), molecular functions (MF) and cellular components (CC), were mainly concerned. g:Profiler is based on Fisher’s one-tailed test, and the statistical *p*-value is multiple testing-corrected. Significant GO terms were filtered by the threshold of “Padj” <0.05. The bar plots of GO enrichment terms were drawn based on R-4.0.3.

### Construction of the weighted gene co-expression network in multi-brain tissues

The fully processed, filtered and normalized gene-level expression profiles from GTEx v8 were used to construct the weighted gene co-expression networks for the top-five brain tissues by R package “WGCNA” (v1.69). WGCNA was performed to build an unsigned gene co-expression network following the standard procedure, and all the parameters were used as recommended, and the soft-threshold was set to 6 after testing a series of soft threshold powers (range 2 to 20). As for the construction of gene co-expression modules, the hierarchical cluster tree in the co-expression network was cut into gene modules using the dynamic tree cut algorithm with a minimum module size of 30 genes^37^. The normalized intra-module connectivity value was computed by setting the options “scaleByMax = T”.

### Drug Gene Interaction (DGIdb) database

DGIdb (v4.2.0) provides a resource of genes that have the potential to be druggable^27^. DGIdb contains two main classes of druggable genome. The first class includes genes with known drug interactions, and the other includes genes that are potentially druggable according to their membership in gene categories associated with druggability. DGIdb includes 42 potentially druggable categories and 49 interaction types (including inhibitors, activators, cofactors, ligands, vaccines and many interactions of unknown types). Only the drug-gene interaction terms with at least one supported PubMed paper were used in the present study.

### PubMed search

To find supports from published research, we performed a text-mining analysis based on PubMed database on June 3rd, 2021. We searched the PubMed database with the items of “((schizophrenia[tiab]+OR+Schizophrenia[tiab]+OR+SCZ[tiab])+AND+(gene name[tiab])+AND+(gene[tiab]+OR+genes[tiab]+OR+mRNA[tiab]+OR+protein[tiab]+OR+proteins[tiab]+OR+transcription[tiab]+OR+transcript[tiab]+OR+transcripts[tiab]+OR+expressed[tiab]+ OR+expression[tiab]+OR+expressions[tiab]+OR+locus[tiab]+OR+loci[tiab]+OR+SNP[tiab]))&d atetype=edat&retmax=100”. The java script output a file with the first column representing gene name, the second column representing the synonyms of the gene name, the last column representing the PubMed ids of hit papers.

### Identification of the potentially phenotype-associated tissues of schizophrenia

To estimate the potentially phenotype-associated tissues, a framework called DESE (also implemented in KGGSEE) proposed by our lab in a recent work was used ^22^. DESE needs three kinds of input datasets, i.e., the expression profiles of various tissues, reference genotype and GWAS summary statistics, and outputs the estimated phenotype-associated tissues.

Specifically, the isoform-level expression profiles of 50 tissues in GTEx v8 were used. The isoform-level expression profiles were preprocessed like this: the index column of the preprocessed expression file was isoform symbol name, and each of 50 tissues or cell types had one column representing the average expression value (i.e., mean value) of corresponding subjects with the tissue. The Genotypes in the EUR panel from the 1000 Genomes Project (phase 3) were downloaded from IGSR and used as reference genotype data. Three columns, i.e., chromosome identifier (CHR), base-pair position (BP) and *p*-value (P) in GWAS summary statistics, were used. SNPs with minor allele frequency (MAF) less than 0.05 were excluded. Only genes approved by HGNC were included in the following analyses. The multiple testing adjustment method was the standard Bonferroni correction, and the cutoff for the adjusted *p*-value was set as p<0.05. The detailed commands of DESE to estimate potential phenotype-associated tissues are described on the KGGSEE website. The bar plot of the rank of potential phenotype-associated tissues was drawn based on R-4.0.3.

### Computation of gene-level eQTLs and isoform-level eQTLs

The present study focused on the cis-eQTLs. Specifically, two files were put into our integrative platform KGGSEE to produce gene/isoform-level eQTLs for each tissue, namely, expression profiles and corresponding genotype data file from GTEx v8. Two levels (gene-level and isoform-level) expression profiles of 50 tissues were downloaded from the GTEx v8 project, and the TMP value was used in the following analyses. Genes/isoforms were selected based on expression thresholds of > 0 TPM in at least 20% of all samples. The genotype data used for eQTL analyses in GTEx release v8 was based on WGS from 838 donors, which all had RNA-seq data available. Only variants with MAF ≥ 0.05 across all 838 samples were included in the present study. GTEx v8 is based on the human reference genome GRCh38/hg38. Thus, to be consistent with the GWAS results of schizophrenia (hg19 assembly), we converted the GRCh38/hg38 coordinates into hg19 by the UCSC LiftOver. All variants were filtered with Hardy–Weinberg disequilibrium (HWD) test *p*-value <1.0E-3. The mapping window was defined as 1 Mb up- and downstream of the gene boundary. If the association test *p*-value of a variant and corresponding expression of gene/isoform was smaller than 0.01, the variant was regarded as a gene-level/isoform-level eQTL of the gene/isoform. It should be noted that the format of the eQTL file is similar to the fasta file. The eQTL data of a gene or isoform starts with the symbol “>”. For the gene-level eQTLs file, the symbol “>” is followed by the gene name (e.g., “*LINC00320*”), its Ensembl ID (“*ENSG00000224924*”) and chromosome identifier (“21”). For the isoform-level eQTLs file, the symbol “>” is followed by the gene name (e.g. “*LINC00320*”), transcript Ensembl ID (“*ENST00000452561*”) and chromosome identifier (“21”). The gene/isoform-level eQTLs files of 50 tissues in GTEx v8 can be accessed on the KGGSEE website and freely used for research purposes. The detailed commands of KGGSEE to compute gene/isoform-level eQTLs of each tissue are described on the KGGSEE website.

### Estimation of the potential susceptibility genes and isoforms for schizophrenia

The framework MCGA included three models, i.e., MCGA_Dist, MCGA_eQTL and MCGA_isoQTL, which were all based on the improved ECS. The main difference among the three models was the strategy used to map variants to genes. For MCGA_Dist, if a variant was within a small window, say +/-5 kb, around the gene boundary, then the variant will be assigned onto the gene according to a gene model, e.g., RefSeqGene. For MCGA_eQTL, the variant will be assigned onto the gene if the variant is a gene-level eQTL of the gene. Similarly, for MCGA_isoQTL, the variant will be assigned onto the isoform if the variant is an isoform-level eQTL of the isoform. Another difference between MCGA_Dist and MCGA_eQTL/MCGA_isoQTL was that the latter two were based on the gene/isoform-level eQTLs of each tissue, thus can produce the potential susceptibility genes/isoforms in a multi-tissue context.

Like our previous model DESE, MCGA contained three iterative steps. In the first step, associated genes with smaller *p*-values of the ECS test were given higher priority to enter the following conditional gene-based association analysis. This step could generate a list of roughly associated genes by removing redundantly associated genes. It should be noted that we dealt with the mentioned three models of MCGA in different ways. For MCGA_Dist and MCGA_eQTL, the order of a gene entering the conditional gene-based association analysis was determined by its *p*-value of the ECS test. For MCGA_isoQTL, assume gene *A* has *m* isoforms. Each isoform could get a *p*-value based on the ECS test, representing the overall statistical significance of all isoform-level eQTLs (simultaneously variants) associated with this isoform. If the isoform with the smallest *p*-value was isoform *a*, with its *p*-value *p_a_*, among the *m* isoforms of gene *A*, we only kept isoform *a* of gene *A* for the following analyses. The adjustment *p*-value for “gene *A* : isoform *a*” pair was adjusted to *m* p_a_* to enter the following conditional gene-based association analysis.

The second step was to compute the selective expression score of genes/isoforms in each tissue by taking all tissues as the background (see details in reference^22^). The Wilcoxon rank-sum test was then performed by using the selective expression score of the associated gene/isoform set and not-associated gene/isoform set (generated by the first step) in each tissue.

In the third step, all genes/isoforms, including the not-associated genes/isoforms, were ranked in descending order based on the tissue-selective expression score of each gene/isoform. The tissue-selective expression score of a gene/isoform was computed based on the rank of this gene/isoform-selective expression score and the *p*-value of the Wilcoxon rank-sum test between the associated gene/isoform set and not-associated gene/isoform set in each tissue.

In the following iteration, genes/isoforms with higher tissue-selective expression scores (in the third step) were given higher priority to enter the conditional gene-based association analysis (in the first step). The above three steps were iterated until the *p*-values of the Wilcoxon rank-sum test did not change almost, and then corresponding associated genes/isoforms were deemed to be potentially associated with the phenotype. More details about the iterative procedure can be found in the original papers^22^.

MCGA is implemented in our integrative platform KGGSEE. To run MCGA_Dist, three input files were needed, i.e., GWAS summary statistics file, gene-level expression profiles of 50 tissues in GTEx v8, genotypes in EUR panel from 1000 Genomes Project (phase 3). To run MCGA_eQTL and MCGA_isoQTL, four input files were needed, i.e., GWAS summary statistics file, gene-level or isoform-level expression profiles of 50 tissues in GTEx v8, genotypes in EUR panel from 1000 Genomes Project (phase 3) and gene/isoform-level eQTLs file of each estimated disease-associated tissue. Only genes with HGNC gene symbols were considered in the present study. The output result file was a text file that contained multiple information about the association measurement of genes (or “gene: isoform” pairs) with the corresponding phenotype. Multiple testing was corrected by using Bonferroni correction. Significant genes were filtered by the “CondiECSp” threshold cutoff 2.5E-6, where “CondiECSp” meant the p-values of conditional gene-based association test based on the improved ECS. The bar plot of the comparison of potential susceptibility genes was drawn based on R-4.0.3. The venn diagram was drawn based on a web app Venny 2.1.0.

### MAGMA

MAGMA is a popular tool for gene and generalized gene-set analysis based on the GWAS summary statistics. Here the parameters and options were used as the MAGMA (v 1.08) manual recommended. Annotation analysis was firstly performed based on the SNP location file and gene location file (hg19, build 37). The SNP location information was extracted from the same GWAS summary statistics file of schizophrenia. An SNP was mapped to a gene if the SNP was in the window of +/-5kb around the gene boundary (same as MCGA_Dist). The gene analysis was performed based on the annotation results and reference data file which was created from Phase 3 of 1000 Genomes of the European population in reference to human genome build 37. Both gene location file and reference data file were downloaded from the MAGMA website. Multiple testing was corrected by using Bonferroni correction. Significant genes were filtered by the threshold of “*p*-value” 2.5E-6.

## Declarations

### Ethics approval and consent to participate

Not applicable.

### Consent for publication

Not applicable.

### Availability of data and materials

The reference genotype data are publicly available in the 1000 Genomes Project^33^ in https://www.internationalgenome.org/. The genotype data are publicly available by application from dbGap (study sccession phg001219.v1) and corresponding gene/isoform-level expression profiles are Genotype Tissue Expression (GTEx v8) project^21^ in https://gtexportal.org/home/. The summary statistics of schizophrenia are publicly available in Psychiatric Genomics Consortium (PGC) in https://www.med.unc.edu/pgc/. The annotations of drug-gene interaction terms are publicly available in Drug Gene Interaction (DGIdb v4.2.0) database^27^ in https://www.dgidb.org/.

The information on FDA-approved antipsychotics can be publicly available in DrugBank 5.0^26^ in https://go.drugbank.com/. The functional enrichment analyses were performed by g:Profiler^36^ and can be publicly available in https://biit.cs.ut.ee/gprofiler. The tool used to draw the venn plot is Venny in https://bioinfogp.cnb.csic.es/tools/venny/index.html. The tool MAGMA^7^ and corresponding reference data were downloaded from https://ctg.cncr.nl/software/magma. Thesource code of MCGA (including MCGA_Dist, MCGA_eQTL and MCGA_isoQTL) is implemented in our integrative software platform KGGSEE and publicly available in http://pmglab.top/kggsee/<u>#/</u>.

## Author’ contributions

M.L. and L.J. conceived the study. M.L. oversaw all aspects of the study. X.L., L.J., C.X, and M.L. developed the models. X.L, L.J. and C.X. performed extensive simulations and real data analyses for performance comparison. X.L., L.J. and M.L. wrote the manuscript. M.L. revised the manuscript. M.J.L. provided useful raw data and did some analysis. All authors commented on and approved the final manuscript.

## Acknowledgments

We thank the GTEx Consortium and 1000 Genomes Projects for providing access to expression and sequence variant data. We also appreciate the authors of the schizophrenia study and the Schizophrenia Working Group of the Psychiatric Genomics Consortium for sharing their GWAS summary statistics.

## Additional Information

### Additional file: Supplemental tables

Table S1: The susceptibility genes of schizophrenia identified by MCGA_Dist.

Table S2: The results of GO enrichment analysis based on the susceptibility genes of schizophrenia identified by MCGA_Dist.

Table S3: The PubMed search hits of the susceptibility genes of schizophrenia identified by MCGA_Dist.

Table S4: The susceptibility genes of schizophrenia identified by MCGA_eQTL.

Table S5: The susceptibility genes of schizophrenia identified by MCGA_isoQTL.

Table S6: The results of GO enrichment analysis based on combined eGene set and combined sGene set of schizophrenia generated by MCGA_eQTL and MCGA_isoQTL.

Table S7: The PubMed search hits of the susceptibility genes of schizophrenia identified by MCGA_eQTL.

Table S8: The PubMed search hits of the susceptibility genes of schizophrenia identified by MCGA_isoQTL.

Table S9: The potential susceptibility isoforms of the twenty-three common genes in the top five phenotype-associated tissues of schizophrenia.

Table S10: The FDA-approved antipsychotics in DGIdb.

Table S11: Drug-gene interaction terms in DGIdb for susceptible genes exclusively predicted MCGA_eQTL.

Table S12: Drug-gene interaction terms in DGIdb for susceptible genes exclusively predicted MCGA_isoQTL.

Table S13: The number of potentially druggable categories for the susceptibility genes of schizophrenia identified by MCGA.

Table S14: The susceptibility genes of schizophrenia identified by MAGMA.

